# Smooth Pursuit Inhibition Reveals Audiovisual Enhancement of Fast Movement Control

**DOI:** 10.1101/2023.09.12.557501

**Authors:** Philipp Kreyenmeier, Ishmam Bhuiyan, Mathew Gian, Hiu Mei Chow, Miriam Spering

## Abstract

The sudden onset of a visual object or event elicits an inhibition of eye movements at latencies approaching the minimum delay of visuomotor conductance in the brain. Typically, information presented via multiple sensory modalities, such as sound and vision, evokes stronger and more robust responses than unisensory information. Whether and how multisensory information affects ultra-short latency oculomotor inhibition is unknown. In two experiments, we investigate smooth pursuit and saccadic inhibition in response to multisensory distractors. Observers tracked a horizontally moving dot and were interrupted by an unpredictable visual, auditory, or audiovisual distractor. Distractors elicited a transient inhibition of pursuit eye velocity and catch-up saccade rate within ∼100 ms of their onset. Audiovisual distractors evoked stronger oculomotor inhibition than visual-or auditory-only distractors, indicating multisensory response enhancement. Multisensory response enhancement magnitudes were equal to the linear sum of responses to component stimuli. These results demonstrate that multisensory information affects eye movements even at ultra-short latencies, establishing a lower time boundary for multisensory-guided behaviour. We conclude that oculomotor circuits must have privileged access to sensory information from multiple modalities, presumably via a fast, subcortical pathway.

## Smooth Pursuit Inhibition Reveals Audiovisual Enhancement of Fast Movement Control

Humans have the remarkable, native ability to react rapidly to novel visual objects or events in their environment. Express saccades can have latencies as short as 80 ms (Fischer & Weber, 1993) and ocular following responses (Miles et al., 1986)—a smooth tracking movement in response to sudden-onset large-field visual motion—can be initiated within 85 ms of motion onset (Gellman et al., 1990). Moreover, the onset of a salient stimulus elicits an involuntary inhibition of microsaccades at ultra-short latencies, a phenomenon known as oculomotor freezing (Engbert et al., 2003; Rolfs et al., 2008l; Hafed & Ignashchenkova, 2013; White & Rolfs, 2016). Such oculomotor inhibition is not limited to saccadic eye movements but has also been observed during smooth pursuit eye movements (Buonocore et al., 2019; Ziv & Bonneh, 2021)—the eyes’ key response to moving objects. In response to a task-irrelevant distractor, ongoing pursuit transiently slows down at a latency of ∼110 ms in humans (Kerzel et al., 2010) and ∼50 ms in monkeys (Buonocore et al., 2019). These studies demonstrate that oculomotor control circuits must have privileged access to sensory signals, allowing the integration of novel environmental events at ultra-short latencies.

Whereas oculomotor inhibition has often been observed in response to visual stimuli, novel events or objects in our natural environment typically contain both light and sound. In general, such cross-modal stimuli trigger faster and more robust neuronal and behavioural responses than stimuli presented via a single sense, a phenomenon known as multisensory response enhancement (e.g., Meredith & Stein, 1983; Rowland et al., 2007; Stein & Stanford, 2008). For instance, pupillary responses are stronger when evoked by audiovisual stimuli as compared to unimodal (visual or auditory) targets (Rigato et al., 2016; Wang et al., 2017; Van der Stoep et al., 2021). Similarly, saccade latencies [Wang et al., 2017; Frens et al., 1995; Corneil et al., 2002; Bell et al., 2005) and manual reaction times are shorter when observers respond to audiovisual targets (e.g., Van der Stoep et al., 2021, Diederich & Colonius, 2004).

Response amplification in the presence of cross-modal stimuli is likely mediated by a number of different brain areas, including the superior colliculus (SC), a midbrain structure that is critically involved in orienting behavior (Allen et al., 2021) including eye movements (Corneil & Munoz, 2014). Intermediate and deep SC layers contain neurons that respond to signals from multiple sensory modalities and exhibit increased firing rates in response to cross-model signals (Meredith & Stein, 1983; Stein & Stanford, 2008). The nature of increased multisensory firing rates can be additive (i.e., multisensory firing rate is equal to the linear sum of the unisensory evoked firing rates), superadditive (i.e., higher firing rate than the linear sum), or sub-additive (i.e., lower firing rate than the linear sum; Stanford et al., 2005; Stanford & Stein, 2007; Stevenson et al., 2014). Many aspects of primate behavior show additive or superadditive multisensory response enhancement.

Here we investigate whether and how cross-modal stimuli affect ultrafast oculomotor inhibition. This is an open question, because (micro)saccadic or pursuit inhibition already occurs at latencies that are at the limit of visuomotor conductance in the brain. Whereas microsaccadic inhibition can exhibit multisensory response enhancement (Wang et al., 2017), it is generally difficult to assess additivity or superadditivity in oculomotor inhibition due to flooring effects. Microsaccade rate, for example, often drops close to zero even in response to unimodal distractors (Rolfs et al., 2008; Wang et al., 2017). One way to overcome this constraint is to investigate pupil responses. However, whereas some studies showed superadditive multisensory pupil responses (Rigato et al., 2016), others observed additive effects (Wang et al., 2017; Van der Stoep et al., 2021). In this study, we use pursuit inhibition to investigate multisensory processing. Pursuit eye movements have two key advantages: (1) in contrast to saccades, which are discrete events, pursuit provides a continuous and gradual response to novel sensory inputs, and (2) compared to pupil orienting, pursuit inhibition occurs faster, providing a fine-tuned window into the temporal evolution of ultrafast sensorimotor responses. Whereas previous studies have established pursuit inhibition in response to unimodal distractors (Kerzel et al., 2010), here we combine signals across both senses to investigate the nature and time course of multisensory inhibition and compare effects on pursuit and catch-up saccades.

## Methods

We conducted two experiments. The study design, sample size and analyses of the main experiment were pre-registered on the Open Science Framework (OSF; https://osf.io/yc3w6). A control experiment was conducted to rule out potential confounds and replicate the main findings at low stimulus contrast. The following study methods apply to the main, pre-registered experiment. Differences between the main and control experiment are described below.

### Observers

We present data from 16 human observers (nine female; 26.6 ± 5.6 years; mean ± SD; four authors) with normal or corrected-to-normal visual acuity and no history of neurological, psychiatric, or eye disease. The sample size was determined using an a priori power analysis (G*Power; Faul et al., 2007) with alpha = .05 and power = .80. Our estimated effect size of *d* = 0.67 was based on pilot data (not included) obtained in *n* = 10 observers. Data from one additional observer were excluded from data analysis because they failed to reliably track the moving target (resulting in >50% excluded trials). The experimental procedure was in accordance with the Declaration of Helsinki and approved by the University of British Columbia Behavioral Research Ethics Board. All observers gave written informed consent and were remunerated at a rate of $10/hour for their participation.

### Apparatus

Observers performed the experiment in a dimly lit laboratory. Stimuli were presented on a gamma-corrected 40.6 cm × 29.8 cm CRT monitor (ViewSonic G255; 85 Hz; 1600 x 1200 pixels) positioned at a viewing distance of 50 cm. An Eyelink 1000 Tower Mount (SR Research, Kanata, ON, Canada) video-based eye tracker recorded observers’ right eye position at a sampling rate of 1 kHz. Observers’ head and chin were supported by a combined forehead and chin rest to minimize head movements. Stimulus presentation and data collection procedures were programmed in MATLAB R2019a (The MathWorks Inc., Natick, MA) using the Eyelink and Psychophysics (version 3.0.12; Cornelissen et al., 2002; Kleiner et al., 2007) toolboxes. Auditory distractors were presented through speakers (EP-691H; SpecResearch, Walnut, CA, USA) placed on either side of the screen (45 cm apart), at a distance of 89 cm from observers.

### Stimuli and procedure

The pursuit target was a small, white Gaussian dot with a standard deviation of 0.14°, presented on a grey background. At the start of each trial, the target was presented 10° to the left of the screen center and then followed a step-ramp motion sequence (**Fig. 1A**): After successful fixation, the target stepped 2° further to the left and then moved rightwards across the screen at a constant velocity of 10°/s for 2,000 ms. Observers were instructed to track the target closely with their eyes until it disappeared from view, marking the end of the trial. In 60% of the trials, a distractor was presented at an unpredictable time within a time window of 500-1,500 ms after ramp onset. Of all distractor trials, 1/3 contained a visual distractor, 1/3 an auditory distractor, and 1/3 a combined audiovisual distractor. The visual distractor was presented for 50 ms and consisted of two horizontally oriented, sinusoidal gratings (1.5 cycles; spatial frequency = 1 cycles per degree; contrast level = 100%), located 5° above and below the target’s horizontal trajectory. A high-contrast, low spatial frequency grating was chosen based on previous studies showing that fast orienting responses are preferentially evoked by high contrast and low spatial frequency stimuli (Ziv & Bonneh, 2021; Kozak et al., 2019). The auditory distractor consisted of a 60 dB white noise sound, presented for 50 ms via both speakers (measured background noise in the laboratory: ∼41 dB). In the audiovisual distractor condition, both distractors were presented simultaneously. In the remaining 40% of trials, no distractor was presented (control condition).

**Figure 1.**
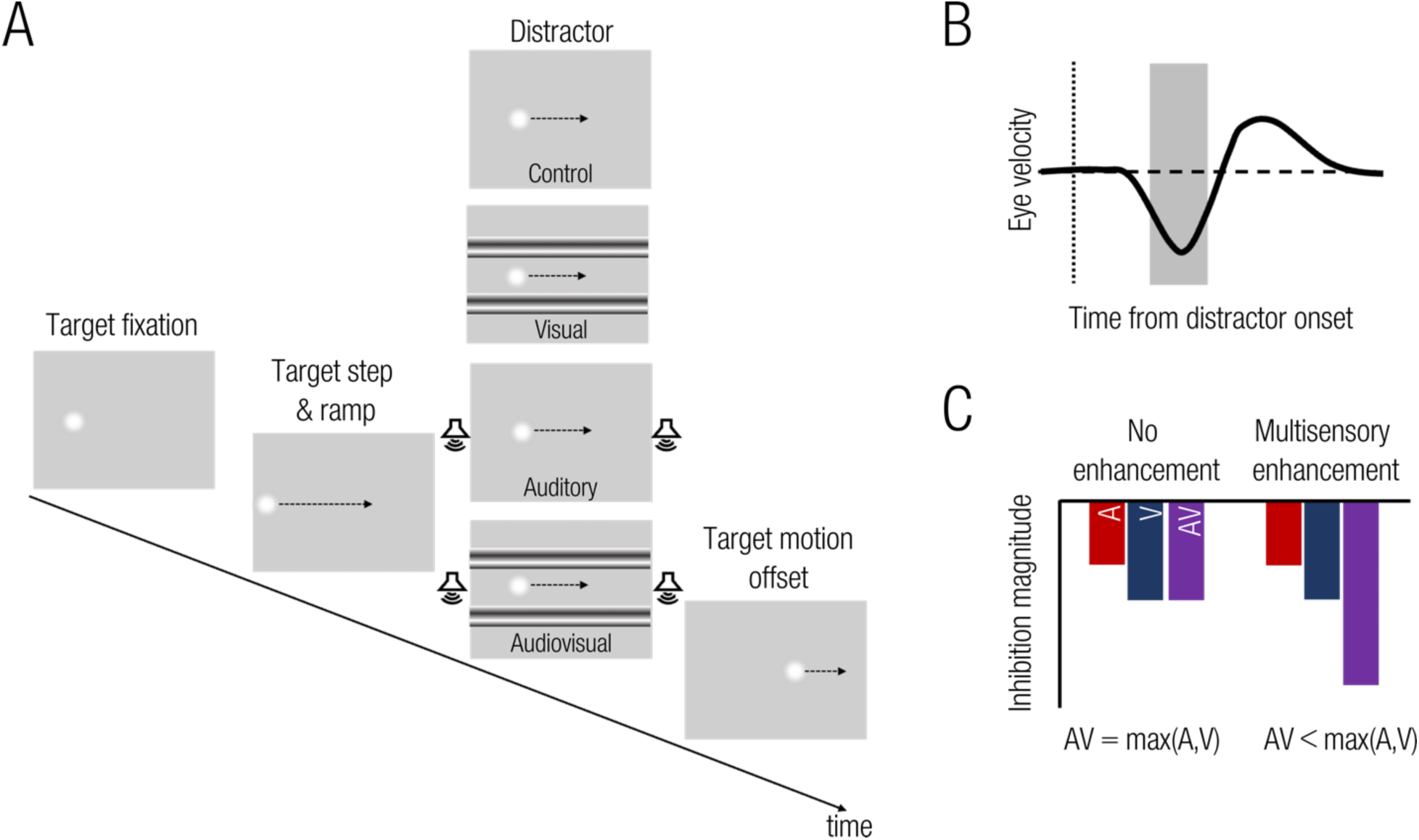
(**A**) Experimental paradigm. Observers tracked a white Gaussian target moving horizontally from left to right. At an unpredictable time (500 to 1500 ms after target onset), a visual (striped pattern), an auditory (white noise), or audiovisual distractor was presented for 30 ms. In the control condition, no distractor was presented. Observers were instructed to track the target until it disappeared, marking the end of the trial. (**B**) Expected inhibition response in pursuit eye velocity evoked by distractor onset (dotted vertical line). An initial, transient inhibition (highlighted grey area) is followed by a rebound period. **(C)** Visualization of the main hypothesis. A: Auditory, V: Visual, AV: Audiovisual.

The experiment was split into 10 blocks of 50 trials each (500 trials in total; 100 in each of the three experimental conditions and 200 in the control condition). Trials were presented in a pseudorandom order. The experiment took approximately 60 minutes to complete.

### Data analyses

*Eye movement data preprocessing.* To investigate the impact of unisensory vs. multisensory distractors on rapid oculomotor inhibition, we analyzed pursuit in a time window from 200 ms before to 400 ms after distractor onset. First, we time-locked all eye movement position traces to the onset of the distractor. For the control condition, we sampled eye movement traces from the same distribution as in the experimental conditions to ensure that eye movement traces from all conditions were sampled over similar time windows. Next, we derived eye velocity traces by digital differentiation of the continuous eye position data. Velocity traces were then filtered using a second-order Butterworth filter with a cut-off frequency of 40 Hz. We detected catch-up saccades using a velocity threshold (30°**/**s), which had to be exceeded for at least five successive frames. The emphasis of our analysis is on the smooth component of the pursuit eye movement, particularly, on pursuit velocity. However, catch-up saccades form an integral part of the pursuit response (Orban de Xivry & Lefèvre, 2007; Goettker & Gegenfurtner, 2021) and we therefore also analyzed the rate of catch-up saccades over time for each observer and experimental condition (see Rolfs et al., 2008; White & Rolfs, 2016). All trials were manually inspected and trials with blinks, undetected saccades, and trials where the eye tracker lost the signal were excluded from further analyses (1.3% of trials).

*Analysis of smooth pursuit and saccade inhibition.* Based on previous studies (Kerzel et al., 2010; Buonocore et al., 2019; Ziv & Bonneh, 2021) we expected the sudden onset of a visual or auditory distractor to elicit a short-latency, transient inhibition of smooth pursuit eye velocity, followed by a rebound period during which pursuit velocity rises above the baseline (**Fig. 1B**). The focus of our analyses is on the latency and magnitude of the early pursuit inhibition response as a measure of multisensory response enhancement. To determine the inhibition onset latency for each experimental condition, we used the jackknife procedure (Miller et al., 1998; Ulrich & Miller, 2001). This method allows accurate detection of small modulations in noisy data. It has been applied to the analysis of event-related potentials (Luck, 2014), modulations in continuous eye movements, such as pupil responses (Grenzbach et al., 2021) or smooth pursuit eye movements (Kerzel et al., 2010). In short, for each condition, we first computed leave-one-out (*n*–1) grand averages. From each grand average, we then determined the maximum dip in eye velocity during the inhibition time window (0 to 200 ms after distractor onset) and defined the inhibition onset latency as the first time point at which eye velocity fell below 50% of the peak inhibition. To account for the reduced variability when basing an analysis on grand averages, we used corrected *F*-values (*F_corr_ = F*/[n–1] ^2^) when statistically comparing latencies between experimental conditions (Luck, 2014). The magnitude of the inhibition response is then defined as the pursuit velocity over a 50-ms time window starting at the mean inhibition onset latency for each observer and condition. Within each experimental condition, we compared the inhibition magnitude to mean baseline pursuit, calculated over a time window from –100 to 0 ms before distractor onset. We applied the same jackknife method to detect inhibition onsets for saccade rates as used for pursuit. A relative criterion of 90% of peak inhibition was used to estimate saccade inhibition magnitude.

### Hypotheses and Statistical Analyses

We analyzed the data in three parts: (1) We confirmed that visual, auditory, and audiovisual distractors elicited a detectable pursuit inhibition response. We compared pursuit baseline and inhibition magnitude using three separate *t*-tests, one for each experimental condition, with a corrected alpha level of .0167. (2) To test whether pursuit inhibition showed a significant multisensory enhancement (**Fig. 1C**), we compared pursuit inhibition magnitudes between the three distractor conditions using a one-way repeated-measures ANOVA (rmANOVA) with factor *distractor type.* We ran Bonferroni-corrected post-hoc comparisons to determine whether the audiovisual distractor elicited significantly stronger inhibition magnitudes as compared to the unisensory (visual or auditory) distractors. For all *t*-tests, we report the Bayes Factor (*BF_10_*), allowing us to assess evidence for the absence of an effect (Keysers et al., 2020). Following established guidelines, we interpret a *BF_10_* of >3 and >10 as moderate and strong evidence for the presence of an effect and a *BF_10_* of <1/3 and <1/10 as moderate and strong evidence for the absence of an effect (Jeffreys, 1961). (3) Multisensory response enhancement is often described as the result of an additive or a superadditive combination of unisensory conditions (Stevenson et al., 2014). We tested these two alternative hypotheses using a linear mixed-effects model on the inhibition magnitudes. We modeled our data as a function of auditory and visual distractor presence or absence, resulting in a 2 (auditory distractor present vs. absent) × 2 (visual distractor present vs. absent) design. If the audiovisual inhibition magnitude can be explained by the linear sum of the unisensory conditions, we would expect significant main effects of visual and auditory distractors, but no significant interaction term. Conversely, if the audiovisual condition produced stronger inhibition than the linear sum of the unisensory conditions, we would additionally expect a significant interaction term. We built the following full model that included both main and interaction effects:

Inhibition Magnitude ∼ Visual Distractor * Auditory Distractor + (1 | observer).

We then compared this full model to a reduced model that only contained the two main effects:

Inhibition Magnitude ∼ Visual Distractor + Auditory Distractor + (1 | observer).

All statistical analyses were performed in R (R Core Team, 2022) with an alpha level of .05 (unless otherwise stated).

*Divergence from the Preregistered Analysis Plan.* According to our initial analysis plan, we aimed to first assess whether the distractors evoked a short-latency inhibition response during smooth pursuit eye movements by comparing inhibition magnitudes between experimental conditions and the control condition using a one-way rmANOVA. We diverged from this plan to account for unequal trial numbers in the control (200 trials per observer) and experimental conditions (100 trials per observer per condition). Additionally, the well-known phenomenon of anticipatory slowing in pursuit velocity toward the end of a trial is problematic in a design that randomizes the onset time of a distractor. To account for this, we first normalized each observers’ pursuit velocity data and then compared inhibition magnitude to a mean baseline pursuit velocity before the distractor onset within each experimental condition (see Kerzel et al., 2010). The originally planned analysis yielded similar results.

### Control Experiment

We recruited an additional 16 observers (eight female; 24.7 ± 3.8 years; mean ± SD; ten observers not tested in the main experiment). Study methods were the same between the main and control experiment with the following exceptions: (1) Visual stimuli were presented on a ViewSonic G90fB CRT monitor (85Hz; 1280×1040 pixels), (2) Visual distractors were presented at three different contrast levels (6.25%, 25%, and 50%), resulting in eight experimental conditions: three visual distractor conditions (each at a different contrast), three audiovisual distractor conditions, one control condition (no distractor), and one auditory distractor condition, presented in pseudorandom order and each repeated 60 times.

## Results

Observers used a combination of smooth pursuit eye movements and catch-up saccades to track the moving target. **Figure 2** shows data from one representative observer, who had a mean pursuit velocity of approximately 10°/s (**Fig. 2A**; left panel), closely matching target velocity, and a catch-up saccade rate of ∼1.2 Hz (**Fig. 2B**; left panel). In this observer, distractors elicited a transient reduction in pursuit velocity and saccade rate ∼100 ms after distractor onset (**Fig. 2**).

**Figure 2.**
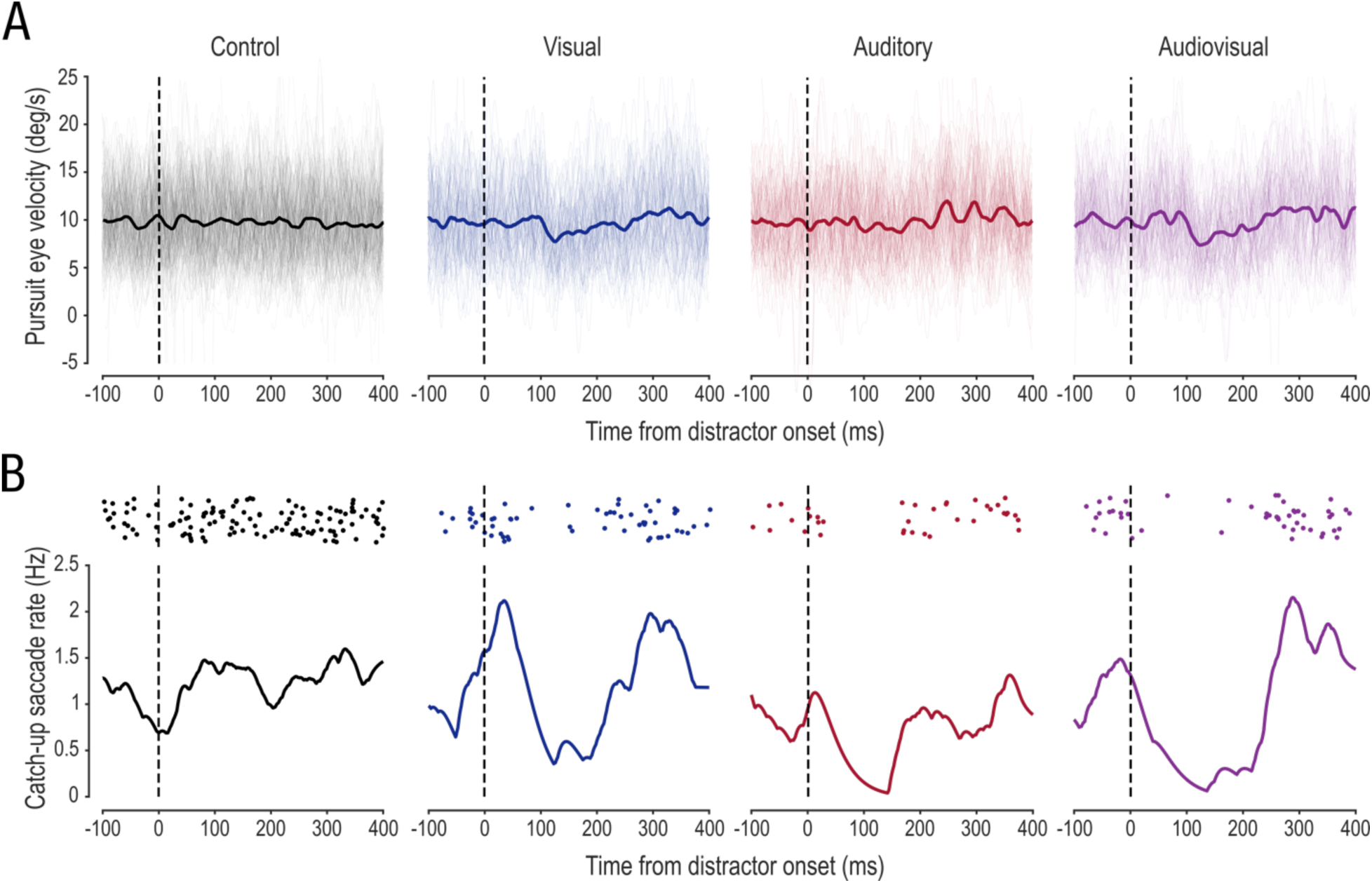
Pursuit velocity and catch-up saccade rates from one representative observer. (**A**) Horizontal smooth pursuit velocity traces from individual trials (light traces) and means across trials (bold traces). (**B**) Raster plots of catch-up saccade onsets (upper panels) and normalized catch-up saccade rates over time (lower panels). Higher overall number of catch-up saccades (upper panel in **B**) in the control condition was due to the higher trial number in this condition. All data are aligned to distractor onset (dashed lines).

### Distractors Elicit Transient Pursuit and Catch-up Saccade Inhibition

Patterns of inhibitory pursuit responses described for the single observer were similar across all observers (**Fig. 3A**). We first confirmed whether different distractors elicited detectable pursuit inhibition responses. To account for individual differences and anticipatory slowing in pursuit, we first normalized pursuit velocity by subtracting the control condition. We then compared mean pursuit velocity during a 50-ms time window from inhibition onset to baseline pursuit in a 100-ms time window before distractor onset. We found a significant reduction in normalized pursuit velocity as compared to baseline pursuit for all three distractor types. The visual distractor elicited a mean velocity reduction of ∼.52°/s (equivalent to a 5.2% reduction in pursuit velocity gain; *t*(15) = –4.23; *p* = .002; *d* = 1.06; *BF_10_* = 50.48). The auditory distractor resulted in a small, yet significant reduction in pursuit velocity of ∼.20°/s (*t*(15) = –3.48; *p* =.01; *d* = .87; *BF_10_* = 13.59). The strongest reduction was observed for the audiovisual distractor, with a mean reduction of ∼.73°/s (*t*(15) = –6.53; *p* < .001; *d* = 1.63; *BF_10_* > 1000; **Fig. 3B**). Thus, all three distractors elicited significant inhibition in pursuit velocity. Moreover, pursuit inhibition responses occurred at short latencies of 93.4 ms to the auditory distractor, 120.5 ms to the visual, and 123.3 ms to the audiovisual distractor (rmANOVA on inhibition latency revealed no differences between distractor types, *F_corr_* < 1).

**Figure 3.**
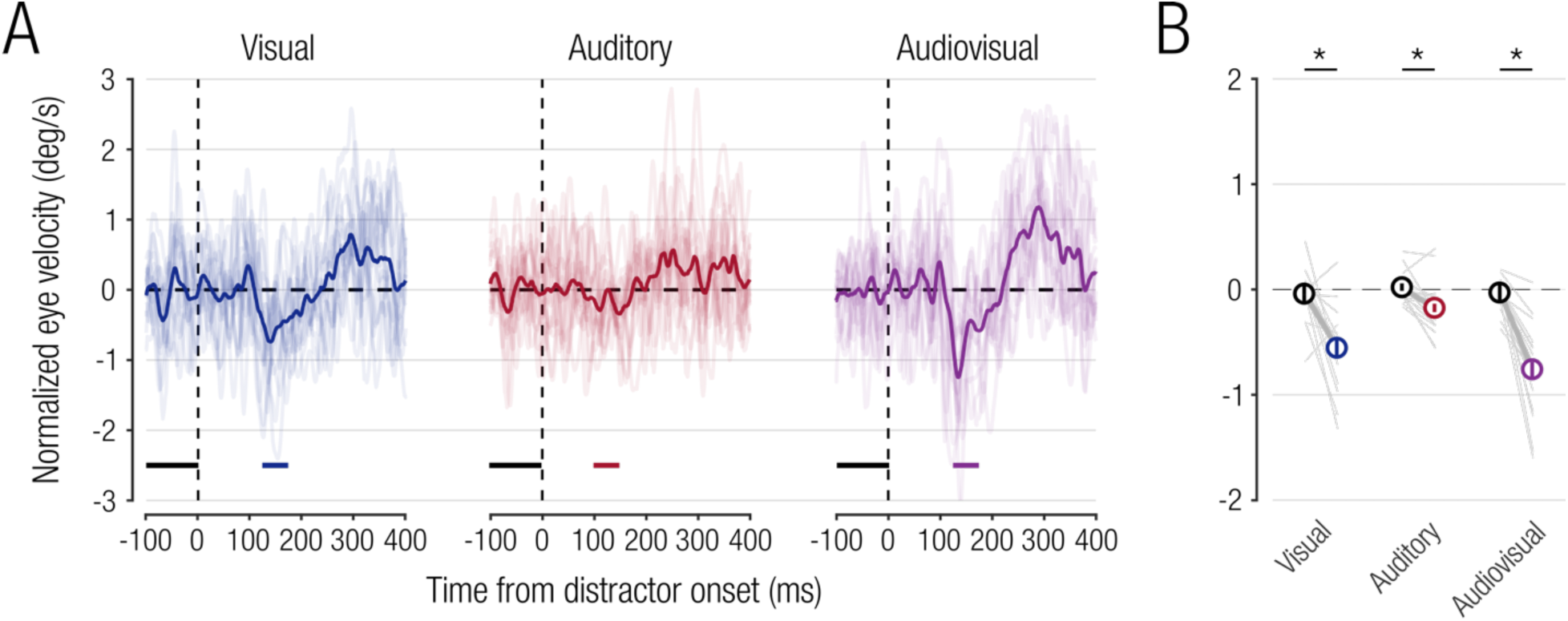
(**A**) Normalized mean pursuit velocity for individual observers (thin lines) and across observers (bold lines). All data were aligned to distractor onset. Horizontal line segments at the bottom of each panel indicate analysis time windows used to calculate baseline pursuit (black) and inhibition magnitudes (coloured). (**B**) Comparison of baseline pursuit and inhibition magnitudes for different distractor types. Circles and errorbars denote mean ± 1 within-subject SEM. Thin grey lines show individual observer data. \**p* < .05.

We next analyzed distractor effects on the catch-up saccade rate as an integral part of the pursuit response (De Brouwer et al., 2002; Orban de Xivry & Lefèvre, 2007; Goettker & Gegenfurtner, 2021). Distractor-induced modulations of catch-up saccade rate followed a similar pattern and time course as observed for smooth pursuit. Distractors elicited a rapid reduction in saccade rate, followed by a rebound (**Fig. 4A**). Saccade rate started to drop at around 70 ms for the visual distractor, and almost immediately after distractor onset for the auditory and audiovisual condition. All three distractors elicited significant reductions in catch-up saccade rates compared to baseline (visual: *t*(15) = –7.18; *p* < .001; *d* = 1.80; *BF_10_* > 1000, auditory: *t*(15) = –6.61; *p* < .001; *d* = 1.65; *BF_10_* > 1000, and audiovisual: *t*(15) = –6.53; *p* < .001; *d* = 1.63; *BF_10_* > 1000; **Fig. 4B**).

**Figure 4.**
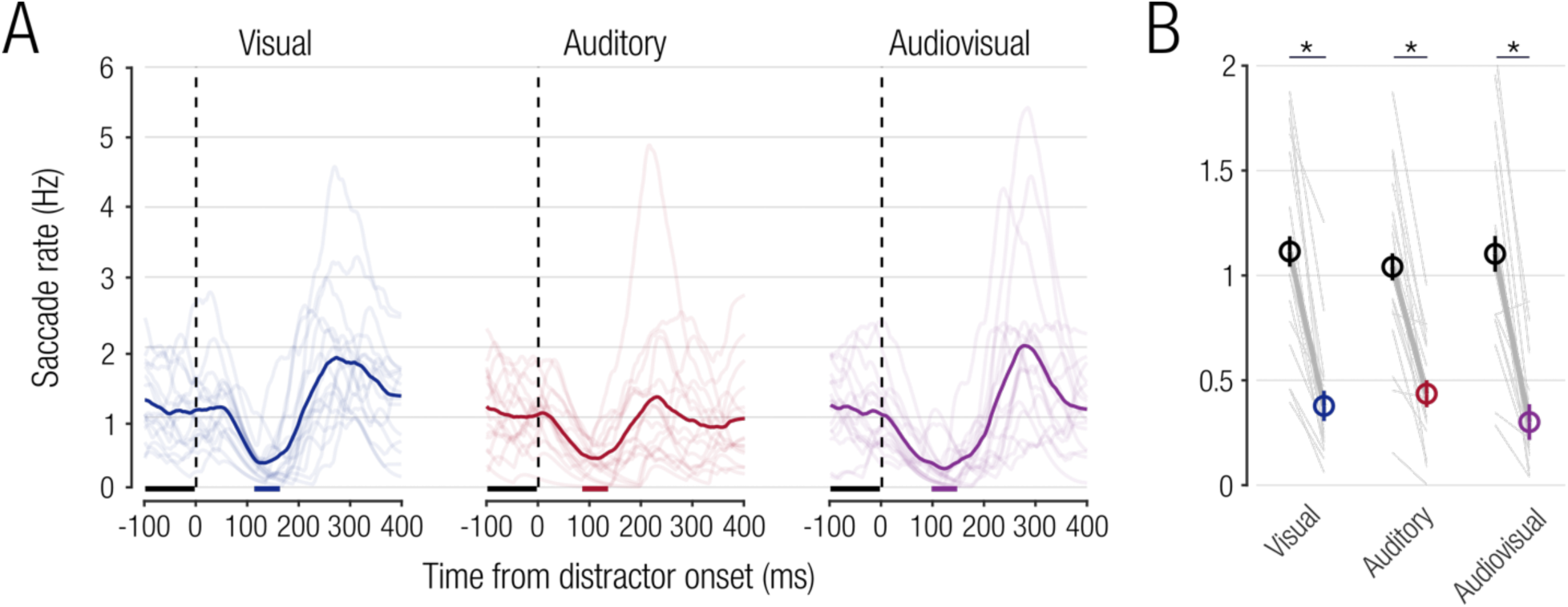
Catch-up saccade rates. (**A**) Individual (thin lines) and mean catch-up saccade rates across 16 observers (bold lines). (**B**) Comparison of baseline catch-up saccade rate and inhibition magnitudes for different distractor types. Same conventions as in fig. 3.

### Multisensory Response Enhancement in Pursuit Inhibition

We next compared pursuit inhibition magnitudes across distractor conditions and asked whether the audiovisual distractor elicited stronger inhibition as compared to the visual or auditory distractors. A one-way rmANOVA yielded a significant main effect of *distractor type* on inhibition magnitudes (*F*(1.47,22.09) = 24.0; *p* < .001; *η_p_^2^* = .74; **Fig. 5A**). Pairwise post-hoc comparisons revealed significantly stronger inhibition in the audiovisual condition as compared to visual (*t*(15) = –2.96; *p* = .019; *d* = .74; *BF_10_* = 5.64; **Fig. 5B**) and auditory distractors alone (*t*(15) = –5.41; *p* < .001; *d* = 1.35; *BF_10_* = 376.73; **Fig. 5C**). **Figures 5B-C** show individual and mean inhibition responses to visual (**Fig. 5B**) and auditory (**Fig. 5C**) distractors compared to the audiovisual distractor. Data points falling along the diagonal indicate similar inhibition responses for unimodal and audiovisual distractors, whereas data points falling below the diagonal indicate stronger inhibition in the audiovisual condition. These results demonstrate that the audiovisual distractor evokes a significantly stronger inhibition effect compared to the unimodal distractors, indicating multisensory response enhancement.

**Figure 5.**
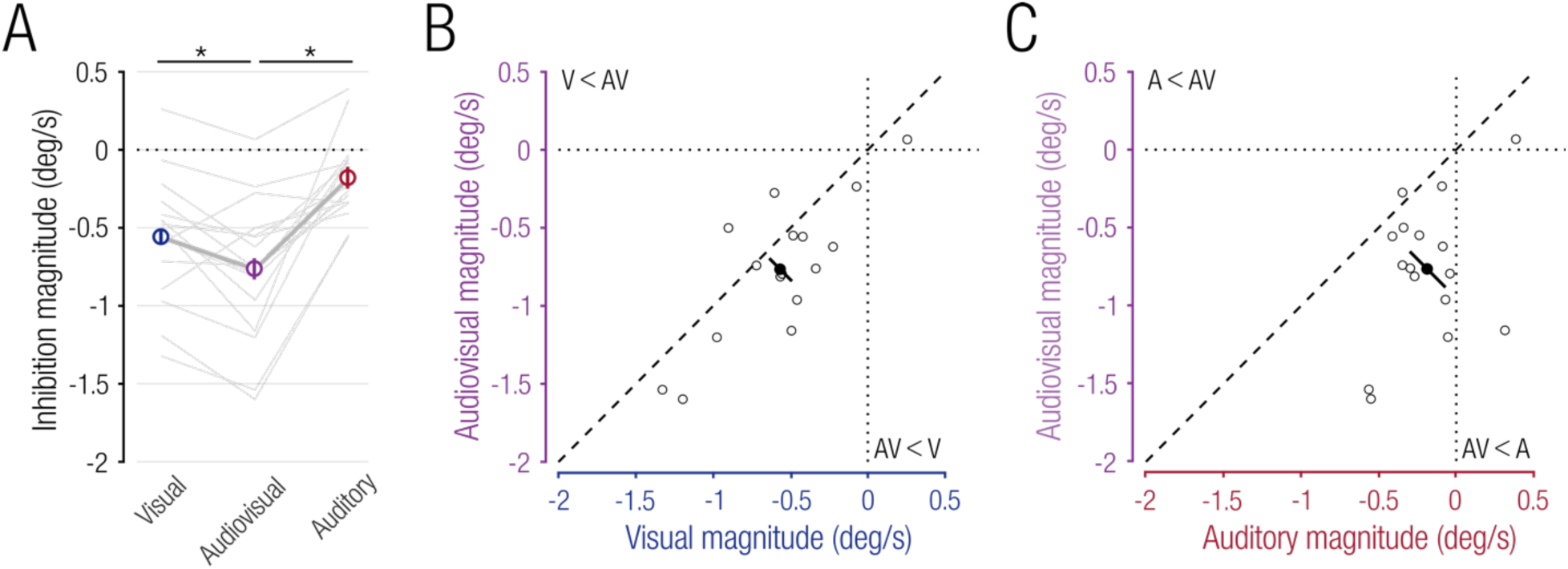
(**A**) Mean inhibition magnitudes. Individual observer trends plotted as thin grey lines. Coloured discs and errorbars represent mean ± 1 within-subject SEM. (**B-C**) Comparison of inhibition magnitudes between unimodal and cross-modal conditions for (**B**) visual and (**C**) auditory distractors. Open circles are individual observer data and filled circles represent the mean across observers. Errorbars show the 95% confidence interval of the mean difference.

Similar to pursuit, the audiovisual distractor also evoked the strongest inhibition of catch-up saccades (**Fig. 4B**). Yet, comparing saccade inhibition magnitudes between visual, auditory, and audiovisual distractors did not reveal a significant effect of *distractor type* on saccade inhibition magnitude (*F*(2,30) = 1.73; *p* = .195). This was likely due to a flooring effect: in several observers, and in contrast to what was observed for pursuit, unimodal distractors evoked a decrease in saccade rate to a value close to zero (e.g., **Fig. 2B**) so that no further response enhancement was possible. The following analyses focus on the pursuit component of the inhibition response.

### Additive Effect of Multisensory Integration in Pursuit Inhibition

Multisensory response enhancement is often attributed to additive or superadditive integration of multisensory signals (Stanford et al., 2005; Stanford & Stein, 2007; Stevenson et al., 2014). We next sought to investigate whether the stronger inhibition response in the audiovisual condition could be explained by an additive or superadditive multisensory effect. To compare these two competing predictions, we analyzed pursuit magnitude as a function of the presence or absence of the visual and auditory distractor. For this analysis, we used mean pursuit velocity from all trials across a 50-ms time window relative to the audiovisual inhibition onset. We ran a linear mixed-effects model with fixed effects *visual distractor* (present vs. absent) and *auditory distractor* (present vs. absent) and random intercepts per observer.

If the multisensory response enhancement resulted from an additive combination of responses in unimodal conditions, we would observe significant main effects of both distractor types but no interaction effect (**Fig. 6A**). Following this logic, the multisensory response can be predicted by adding the effects of the visual distractor (baseline vs. visual distractor only) and auditory distractor (baseline vs. auditory distractor only). Conversely, a superadditive effect would predict a significant interaction term between the two distractor types (i.e., stronger effect if both distractors are present compared to the linear sum of both distractors; **Fig. 6A**). The full model revealed main effects of the visual (*F*(1,7731) = 133.06; *p* < .001) and auditory distractors (*F*(1,7731) = 15.56; *p* < .001) and no interaction effect (*F*(1,7731) = .11; *p* = .741; **Fig. 6B**).

**Figure 6.**
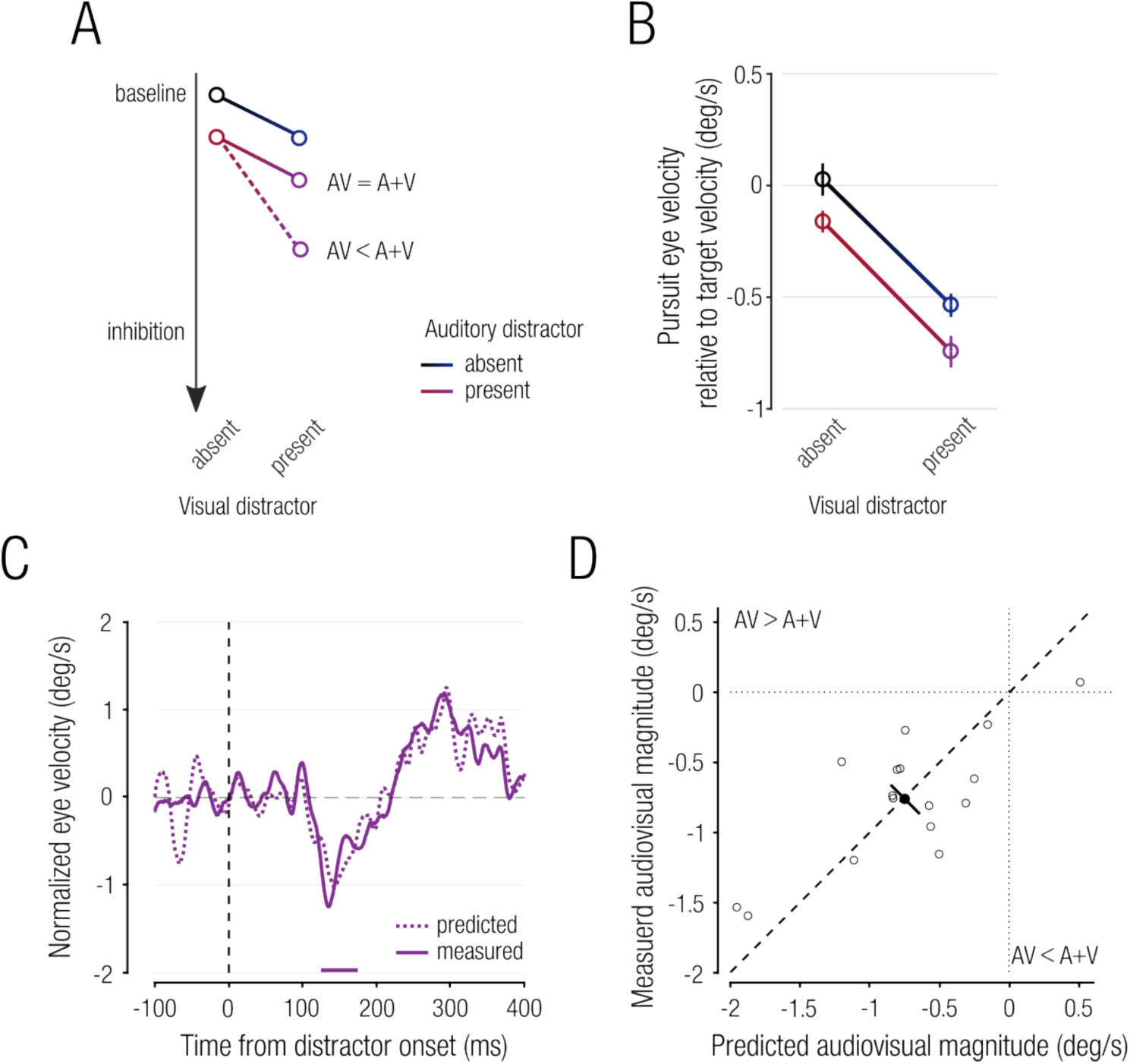
(**A**) Predictions of additive (AV = A+V) and superadditive (AV < A+V) effects of multisensory response enhancement on pursuit inhibition. (**B**) Observed inhibition responses as a function of the presence or absence of the visual and auditory distractors. (**C**) Mean measured normalized pursuit velocity in the audiovisual condition (solid line) and mean predicted pursuit velocity based on the linear sum of the measured unisensory conditions (dashed line). (**D**) Comparison of predicted and measured inhibition magnitudes. Open circles are individual observer data and filled circles represent the mean across observers. Errorbars show the 95% confidence interval of the mean difference.

Comparing the full model to a reduced model without the interaction term did not yield a significant difference between models (*χ^2^*(1) = .11; *p* = .741; AIC for full model: 33676 versus reduced model: 33674), indicating that the observed multisensory response enhancement was not superadditive. To test the alternative that the multisensory response was additive, we used data from the unimodal conditions to predict an additive response. Indeed, a comparison of the predicted (dashed line in **Fig. 6C**) and measured audiovisual responses (solid line in **Fig. 6C**) revealed no significant difference (*t*(15) = .18; *p* = .860; *d* = .04; *BF_10_* = .26; **Fig. 6D**). A Bayes Factor of < .33 indicates moderate evidence in favor of the null hypothesis that there is no difference between the predicted and measured inhibition magnitudes. Therefore, the enhanced pursuit inhibition caused by the audiovisual distractor can be well explained by an additive combination of the component stimuli.

### Additive Signal Integration at Low Distractor Contrast

The multisensory response enhancement observed in pursuit inhibition magnitude can be explained by an additive effect. Similar findings have been obtained with pupil responses to multisensory stimuli (Van der Stoep et al., 2021). Van der Stoep and colleagues’ study and our main experiment used high-contrast visual stimuli, potentially causing a flooring effect in pursuit inhibition (or pupil responses).

Therefore, to assess whether additive effects of multisensory response enhancement hold for low-intensity stimuli, we manipulated visual distractor contrast in a control experiment. Observers performed the same task as in the main experiment, but with the visual distractor presented at either 6.25%, 25%, or 50% contrast. Similar to the main experiment, we again observed a transient distractor-induced inhibition in pursuit eye velocity, subsequently followed by a rebound (**Fig. 7**). First, we confirmed that the lower visual contrasts in the control experiment evoked smaller pursuit inhibition magnitudes in the audiovisual distractor condition, as compared to the main experiment. Using Welch *t*-tests with a corrected alpha level of .0167, we found significantly smaller inhibition magnitudes in the 6.25% (*t*(29.98) = –2.68; *p* = .035; *d* = .95; *BF_10_* = 4.49) and the 25% contrast conditions (*t*(28.74) = –2.88; *p* = .022; *d* = 1.02; *BF_10_* = 6.47). There was no significant difference between the 50% contrast condition as compared to data obtained in the main experiment (*t*(28.64) = –1.39; *p* = .525; *d* = .49; *BF_10_* = .70).

**Figure 7.**
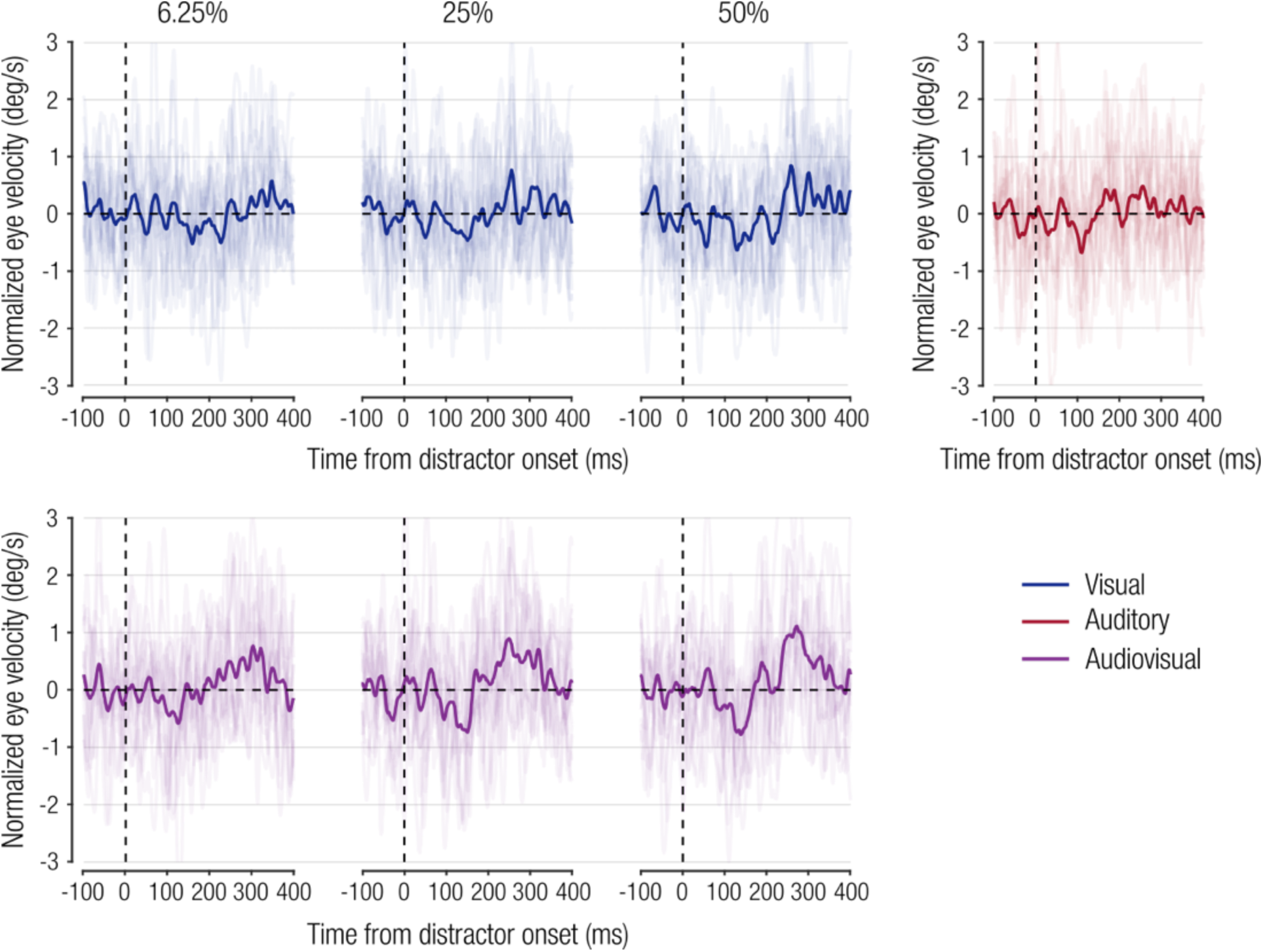
Distractor effect on pursuit eye velocity in the control experiment using different visual contrast levels (6.25%, 25%, and 50%). Thin lines represent individual pursuit velocity traces and bold lines represent means across observers. Note, the lower number of trials per condition in the control as compared to the main experiment resulted in overall noisier pursuit eye velocity traces.

We next assessed whether an additive combination of auditory and visual distractors predicted responses evoked by audiovisual distractors, akin to the main experiment. Predicted responses based on additive combination of the component stimuli (dashed lines in **Fig. 8A**) closely resembled the measured audiovisual condition (solid lines in **Fig. 8A**). Congruently, we found no differences between predicted and observed responses for any of the three contrast levels and Bayes Factors imply moderate evidence for the absence of an effect (6.25% contrast: *t*(15) = .31; *p* = 1; *d* = .08; *BF_10_* = .27; 25% contrast: *t*(15) = .94; *p* = 1; *d* = .24; *BF_10_* = .38; 50% contrast: *t*(15) = –.11; *p* = 1; *d* = .03; *BF_10_* = .26; **Fig. 8B**). These findings indicate that an additive combination of auditory and visual distractors could explain pursuit inhibition to audiovisual distractors, irrespective of contrast levels.

**Figure 8.**
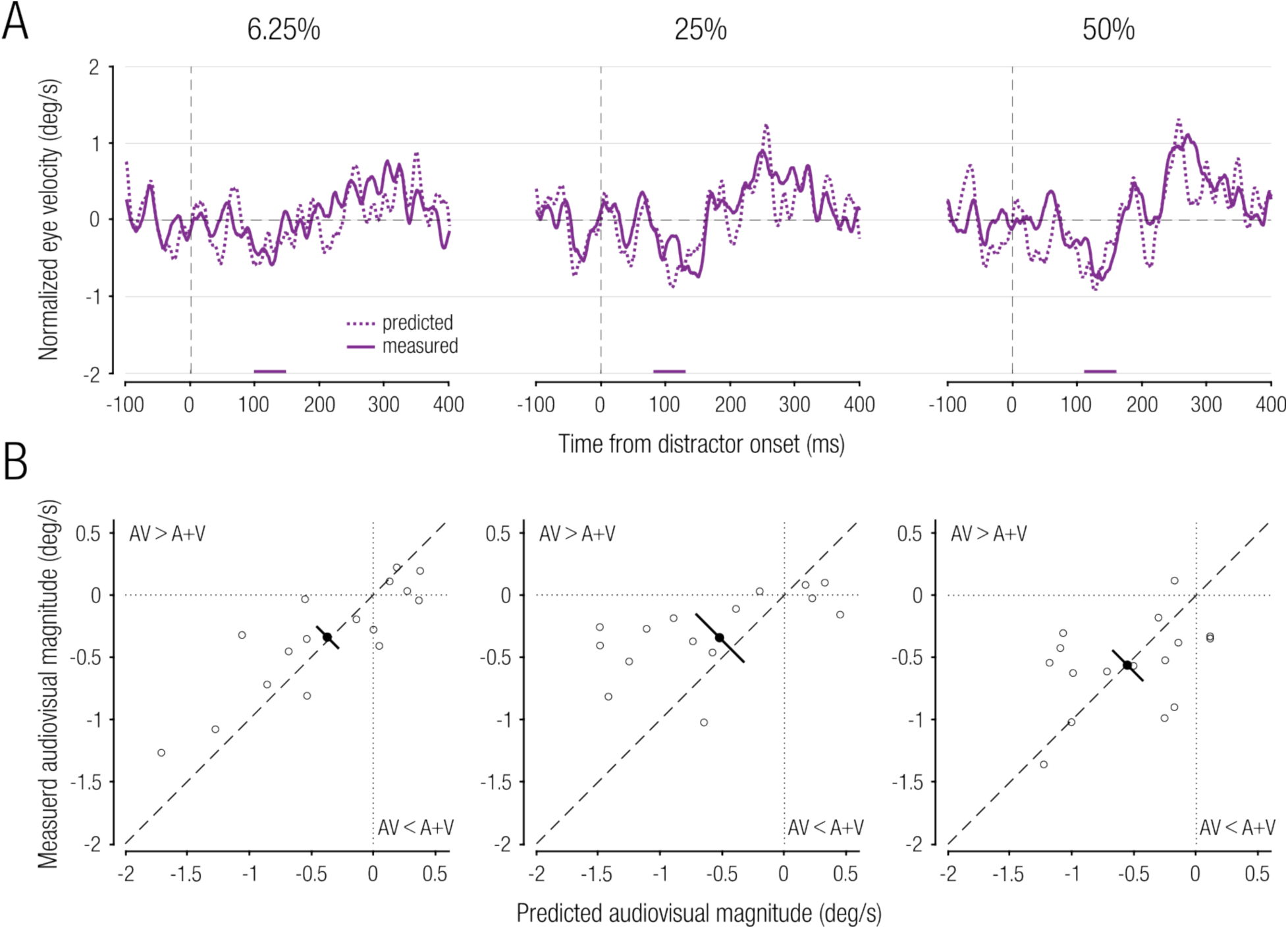
(A) Measured normalized pursuit velocity in response to the audiovisual distractors (solid lines) and predicted normalized pursuit velocity based on the linear sum of the measured unisensory conditions (dashed lines). **(B)** Comparison of predicted and measured inhibition magnitudes. Open circles show individual data and filled circles represent means across observers. Errorbars show 95% CI of the mean difference between conditions. Note, the noisier pursuit velocity traces were due to the lower number of trials per condition in the control experiment.

## Discussion

Oculomotor inhibition in response to sudden-onset distractors occurs at ultra-short latencies and provides a behavioural measure of rapid visuomotor processes in the human brain. We used smooth pursuit eye velocity and catch-up saccade rate to probe oculomotor inhibition in response to visual, auditory, or audiovisual distractors. Distractor onsets elicited a rapid decrease in pursuit eye velocity and catch-up saccade rate. Critically, when probed with synchronous audiovisual distractors, pursuit inhibition was significantly stronger as compared to either visual or auditory distractors alone, indicating multisensory response enhancement. An oculomotor orienting response that is enhanced by cross-modal stimulation might be an advantageous mechanism to increase an organism’s efficiency in detecting and discriminating between sudden external events in the environment.

Whereas saccades are discrete events that are often completely inhibited in response to either visual or auditory distractors, smooth pursuit eye movements are continuous and provide a gradual measurement of distractor-evoked oculomotor inhibition. Pursuit inhibition allowed us to explore the mechanism of multisensory processing for oculomotor inhibition. We show that the multisensory response enhancement magnitude was equal to the linear sum of the responses evoked by the component stimuli. This additive signal combination was observed across visual contrast levels, indicating that additive multisensory response enhancement affects oculomotor inhibition across suprathreshold contrast levels.

Our findings are in line with previous studies showing effects of multisensory response enhancement on the latency of saccades (e.g., Corneil et al., 2002; Bell et al., 2005; Wang et al., 2017) or the latency and magnitude of pupil responses (Rigato et al., 2016; Wang et al., 2017; Van der Stoep et al., 2021). Wang and colleagues (2017) observed significantly stronger microsaccadic inhibition to audiovisual stimuli, as compared to visual or auditory stimuli, but this study did not distinguish between additive and superadditive multisensory enhancement. Here we provide new evidence for an additive effect of multisensory response enhancement for rapid oculomotor inhibition. Moreover, our results show that cross-modal stimuli affect eye movements at very short latencies, a finding that would be more difficult to establish with discrete behaviours such as saccades and button presses, which typically have latencies of more than 200 ms. These results suggest that information from the two senses might converge faster and earlier to control behaviour than previously thought.

### Additive Multisensory Integration in Orienting Behavior

Both additive and superadditive multisensory responses indicate the use of signals across modalities. Yet, some consider only superadditive effects conclusive behavioural evidence for the actual integration of signals due to involvement of non-linear signal combinations that cannot be explained by independent sensory processes alone (Stevenson et al., 2014). Following this logic, additive effects could also result from statistical facilitation of two separate sensory processes and might not necessarily require multisensory convergence and integration at the single neuron level. Human behavioural and neurophysiological studies on multisensory response enhancement have therefore focused primarily on superadditive combinations of multisensory stimuli. This emphasis on superadditive effects was initially supported by single-cell recordings showing firing rates to cross-modal stimuli that far exceed the linear sum of neural activity caused by the component stimuli (Meredith & Stein, 1983). However, subsequent studies demonstrate that superadditive multisensory enhancement is primarily observed when combining subthreshold or near-threshold unimodal stimuli (Stanford et al., 2005). With increasing stimulus strength, most neurons exhibit firing rates that approximate the linear sum of unimodal influences (Stanford & Stein, 2007).

These findings suggest that superadditivity is only a special case, rather than a hallmark, of multisensory integration that occurs when probed with near-threshold stimuli. In fact, the strict focus on superadditivity has been challenged (Stanford & Stein, 2007; Angelaki et al., 2009). Additive and even subadditive operations are common neural mechanisms of multisensory integration when tested with suprathreshold stimuli (Angelaki et al., 2009). Although we did not systematically assess observers’ perceptual thresholds, all stimuli used in our experiments were designed to be well above perceptual thresholds. Our findings of additive multisensory enhancement are therefore in line with the idea of additive multisensory integration when probed with suprathreshold stimuli. Future studies could use a wider range of stimulus strength, including near and subthreshold unimodal stimuli, to probe whether stronger multisensory enhancement is observed when using weaker stimuli.

### Common Mechanism for Pursuit and Saccade Inhibition?

Our results, together with the results of two previous studies (Buonocore et al., 2019; Ziv & Bonneh, 2021), suggest a common inhibitory mechanism for pursuit and saccades, causing the eyes to rapidly freeze, or slow down, in response to the sudden onset of a salient event. Our results extend these findings showing that pursuit and saccade inhibition are stronger in response to cross-modal, as compared to unimodal stimuli.

Due to their short latency, pursuit and (micro-)saccadic inhibition may be mediated by a fast, subcortical pathway (Hafed et al., 2021). Previous studies suggested a critical role of the superior colliculus (SC) in mediating (micro)saccadic inhibition (Engbert, 2006). The SC also plays a key role in the integration of cross-modal signals (Meredith & Stein, 1983; Stein & Stanford, 2008) and the coordination of various aspects of the oculomotor orienting response, such as head movements, saccades, and pupil response (Sparks, 1986; Corneil & Munoz, 2014). Although smooth pursuit is primarily driven by sensory signals in motion-sensitive middle temporal visual area (Lisberger & Movshon, 1999), the SC also encodes visual saliency during smooth pursuit eye movements (White et al., 2021), making it a candidate to mediate distractor-induced pursuit modulations. Recent neurophysiological findings question a causal role of the SC in mediating oculomotor inhibition and instead point to omnipause neurons (OPN) in the brainstem (Hafed et al., 2021). Tonically firing OPNs prevent unintended saccades by inhibiting premotor saccadic burst neurons (Sparks, 2002). The same inhibitory mechanism has also been suggested for smooth pursuit eye movements (Missal & Keller, 2002; Krauzlis, 2004). Therefore, OPNs are another possible candidate mediating inhibitory effects observed across different types of eye movements. However, whether and how OPNs respond to cross-modal stimuli is currently unknown.

Whereas our results and other behavioural and neurophysiological studies suggest a common mechanism controlling pursuit and saccadic inhibition, we also observed some noteworthy differences. First, the saccade inhibition was larger than pursuit inhibition: peak pursuit inhibition was ∼10% of baseline pursuit velocity, but saccade rates decreased by ∼70% and even dropped to zero in some observers. Second, auditory distractors elicited weak pursuit inhibition (**Fig. 3**) but strong inhibition of catch-up saccade rates (**Fig. 4**). This difference indicates that pursuit eye movements might be less affected by auditory signals (Kerzel et al., 2010), congruent with the observation that humans have limited ability to smoothly track non-visual motion signals (Berryhill et al., 2006).

## Conclusion

Combining sensory inputs from different modalities is a vital capability resulting in increased response magnitudes, decreased reaction times, and lower detection thresholds, enabling more efficient reactions to a changing environment. The ability to rapidly detect and respond to changes extends to distractors—interfering and task-irrelevant events that must be suppressed. Here we provide new evidence for an additive effect of cross-modal signals on the magnitude of short-latency smooth pursuit inhibition, occurring reflexively to sudden-onset distractors. Due to the continuous nature of pursuit, its inhibition can provide a testbed to examine distractor suppression (Spering et al., 2006; Wöstmann et al., 2022) and a trial-by-trial window into short-latency multisensory processing.

## Acknowledgements

The authors would like to thank Alex L. White for help with the catch-up saccade rate analyses and members of the Spering lab for comments on an earlier version of the manuscript.

